# Nonalcoholic fatty liver disease is associated with decreased hepatocyte mitochondrial respiration, but not mitochondrial number

**DOI:** 10.1101/2020.03.10.985200

**Authors:** MC Sinton, J Meseguer Ripolles, B Lucendo Villarin, MJ Lyall, RN Carter, NM Morton, DC Hay, AJ Drake

**Affiliations:** University/BHF Centre for Cardiovascular Science, Queen’s Medical Research Institute, Edinburgh BioQuarter, 47 Little France Crescent, Edinburgh EH16 4TJ; Scottish Centre for Regenerative Medicine, Edinburgh BioQuarter, 5 Little France Crescent, Edinburgh, EH16 4UU

**Author notes:** **Corresponding author:** Amanda J. Drake: University/BHF Centre for Cardiovascular Science, Queen’s Medical Research Institute, Edinburgh BioQuarter, 47 Little France Crescent, Edinburgh EH16 4TJ.

## Abstract

Nonalcoholic fatty liver disease (NAFLD) is currently the most prevalent form of liver disease worldwide. This term covers a spectrum of pathologies, from benign hepatic steatosis to non-alcoholic steatohepatitis (NASH). As the disease progresses, NASH can develop into cirrhosis and hepatocellular carcinoma. However, the underlying mechanisms and the factors which predispose an individual to disease progression remain poorly understood. Whilst NAFLD appears to be associated with mitochondrial dysfunction, it is unclear whether this is due to respiratory impairment, changes in mitochondrial mass, or mitochondrial fragmentation. Using a human pluripotent stem cell-based model of NAFLD we show that exposure to lactate, pyruvate and octanoic acid results in the development of macrovesicular steatosis. We do not observe changes in mitochondrial mass or fragmentation but do find decreases in maximal respiration and reserve capacity, suggesting impairment in the electron transport chain (ETC). Taken together, these findings indicate that the development of macrovesicular steatosis in NAFLD may be linked to the impairment of the ETC in mitochondria.

## Introduction

Non-alcoholic fatty liver disease (NAFLD) is the most prevalent form of hepatic disease globally, estimated to affect 1 in 3 people. It is closely related to obesity, with ∼88% of obese individuals at risk of developing the disease [1]. NAFLD is characterised by excess accumulation of triglycerides (TGs) in hepatocytes, leading to the development of steatosis [2]. Steatotic hepatocytes accumulate TGs in single large, or multiple medium-sized, cytoplasmic lipid droplets [3] and whilst simple steatosis is largely benign, its presence increases the risk of developing non-alcoholic steatohepatitis (NASH), which may progress to cirrhosis and hepatocellular carcinoma [4]. It is unclear why only a subset of patients develop NASH and, other than bariatric surgery for morbidly obese patients, there are currently no specific therapeutics available to treat or reverse this pathology [5].

Multiple studies suggest that the pathology of NAFLD is linked to impairment of mitochondrial function[6–9]. In rat models, mitochondrial dysfunction precedes the development of hepatic steatosis, suggesting that disease pathology and progression are linked to impairment of mitochondrial function [10]. NAFLD is linked to structural changes in mitochondrial cristae and the development of crystalline inclusion bodies [11]. As electron transport chain (ETC) components are embedded in the cristae, structural changes may have a physical impact on respiration [12]. Similarly, the growth of inclusion bodies may also impair oxidative phosphorylation and lead to increased free radical production [11]. This is supported by observations of NAFLD-associated hepatic oxidative stress and altered levels of fatty acid beta oxidation [11,12]. This is supported by the observation of NAFLD-associated hepatic oxidative stress and altered levels of fatty acid beta oxidation [11,12]. Furthermore, in humans, once hepatic lipid content exceeds 10%, lipid export is compromised [13]. In tandem with altered fatty acid beta oxidation, this imbalance may lead to the accumulation of intracellular lipids observed in NAFLD.

Satapati et al. (2012) observed increased tricarboxylic acid (TCA) cycle activity and anaplerosis in the livers of mice with high fat diet-induced hepatic steatosis [7]. They also demonstrate that increased levels of hepatic TGs induced oxidative metabolism, with a proportional increase in oxidative stress [14]. Indirect studies in humans indicate a similar effect on TCA cycle flux [6]. As TCA cycle flux generates NADH and FADH_2_, which transfer electrons to the electron transport chain (ETC), alterations in TCA cycling may have a direct impact on respiration and oxidative stress. Further studies in humans demonstrate that NAFLD and NASH are associated with increased and decreased mitochondrial maximal respiration, respectively, indicating that there is a transition in mitochondrial function between these two disease states [15]. The reasons for this are unclear; such differences in maximal respiration between NAFLD and NASH could reflect impairment of the ETC and/or changes in baseline mitochondrial respiration. Furthermore, mitochondria isolated from whole tissue come from multiple cell types, which may differ in proportion in disease states and this may confound the interpretation of any results. Observations that mitochondrial mass increased in NASH with a concomitant decrease in maximal respiration may also arise from organelle fragmentation, a phenotype associated with oxidative stress and apoptosis in other diseases [16].

In light of current knowledge, we hypothesised that nutrient excess in NAFLD leads to mitochondrial dysfunction and subsequent increases in ETC activity due to increased availability of respiratory substrates. Therefore, we aimed to determine the impact of lipid accumulation on mitochondrial respiration specifically in hepatocytes, to avoid the confounding effects associated with bulk tissue analysis. We also aimed to determine whether changes in respiration were related to altered mitochondrial mass or fragmentation. To this end, we used a recently developed *in vitro* model of NAFLD [17] to explore the impact of hepatic steatosis on mitochondrial respiration.

## Methods

### Differentiation of pluripotent human stem cells to hepatocyte-like cells and induction of intracellular lipid accumulation

Human female H9 pluripotent stem cells (PSCs) were differentiated to hepatocyte-like cells (HLCs) as previously described [18]. HLCs were cultured in a 96-well format for measurements of lipid accumulation and in a 6-well format for all other analyses. Intracellular lipid accumulation was induced in HLCs, as previously described [17]. Briefly, at day 17, HLCs were exposed to a cocktail of sodium l-lactate (L; 10mM), sodium pyruvate (P; 1 mM) and octanoic acid (O; 2 mM) (Sigma, Gillingham, UK).

### High content analysis microscopy

Cells were stained with a cell painter assay, adapted from Lyall *et al* and Bray *et al* [17,19]. Cells were fixed with 50 μL/well 4% (wt/vol) paraformaldehyde (Electron Microscopy Sciences, 15710-S) for 15 minutes at room temperature. For permeabilisation, cells were incubated in 0.1% Triton X-100 (Sigma-Aldrich, T8787) in PBS for 15 minutes at room temperature. For lipid droplet analysis, cells were then stained with a combination of NucBlue Live ReadyProbes^®^ Reagent (2 drops/mL) (Molecular Probes, R37605), HCS CellMask™ Red (2 μL/10 mL) (Invitrogen, H32712), and BODIPY™ 493/503 (1:1000) (Life Sciences, D3922), as per the manufacturer’s instructions. For mitochondrial quantification, cells were stained with a combination of NucBlue Live ReadyProbes^®^ Reagent (2 drops/mL), MitoTracker™ Deep Red FM (1/4000; Invitrogen, M22426), and Alexa Fluor™ 555 Phalloidin (Invitrogen, A34055). Following staining, images were acquired using an Operetta High Content Analysis microscope (Perkin Elmer, Buckinghamshire, UK). Lipid droplet morphology was analysed as previously described [17].

### Cell mitochondrial stress test assay

The oxygen consumption rate (OCR) of LPO-exposed HLCs was measured using the Agilent Seahorse XF Cell Mito Stress Test Kit (Agilent, 103015-100) on a Seahorse XF Analyser (Agilent, California, USA). Analysis was performed under basal conditions, and following treatment with oligomycin A (an ATPase inhibitor), carbonyl-cyanide-4-(trifluoromethoxy) phenylhydrazone (FCCP; an ETC uncoupler), and combined rotenone and antimycin A (inhibitors of complex I and III, respectively). Two concentrations of FCCP (0.5 μM and 1.0 μM) were used for optimisation. Since replicates within each group responded similarly to each other, results were combined. OCR was normalised to total protein for each well, using the sulforhodamine B (SRB) assay, as previously described [20], but with spectrophotometric measurements read at 540 nm.

### Citrate synthase assay

Mitochondria were isolated using the Mitochondria Isolation Kit for Cultured Cells (Thermo Scientific, 89874), as per the manufacturer’s instructions, selecting option A for isolation. Citrate synthase activity, a marker for mitochondrial integrity, was then measured using the Citrate Synthase Activity Colorimetric Assay Kit (BioVision, K318), as per the manufacturer’s instructions.

### Protein Extraction

Adherent HLCs were washed once with ice-cold PBS, before incubating in ice-cold RIPA Lysis and Extraction Buffer (Thermo Scientific, 89900) supplemented with cOmplete™ Protease Inhibitor Cocktail tablets (1/10 mL buffer; Roche, 11697498001). The suspended HLCs were placed on ice for 30 minutes, vortexing every 3 minutes, before centrifuging for 20 min at 4 °C, 12,000 rpm. The supernatant was collected and stored at −80 °C until needed.

### Western blot analysis

Protein quantification was performed using the Qubit™ Protein Assay Kit (Invitrogen, Q33211), as per the manufacturer’s instructions. Protein concentration was measured using a Qubit™ Fluorometer (Invitrogen, Massachusetts, USA). Equal concentrations (50 μg) of HLC protein extract in 4 × Sample Loading Buffer (Li-Cor, 928-40004) were loaded onto NuPAGE™ 4-12% Bis-Tris Protein Gels (Invitrogen, NP0326BOX). Following resolution, protein was transferred to a methanol-activated polyvinylidene difluoride (PVDF) membrane. Protein transfer was measured using Revert 700 Total Protein Stain Kit (Li-Cor, 926-11010) as per the manufacturer’s instructions. Membranes were then blocked with Tris-buffered saline containing Tween 20 (TBST) and 5% skimmed milk powder, before incubating with either Pyruvate Dehydrogenase (C54G1) Rabbit mAb (Cell Signaling Technology, 3205) or SDHA (D6J9M) XP^®^ Rabbit mAb (Cell Signaling Technology, 11998), both a 1:1000 dilution. The membranes were washed in TBST before incubating with the secondary antibody, IRDye^®^ 680RD Donkey anti-Mouse IgG (Li-Cor, 926-68072) at a 1:10,000 dilution, for 1 h at room temperature, in the dark, with shaking. Blots were visualised on a Li-Cor Odyssey^®^ CLx (Li-Cor, Nebraska, USA), and bands normalised to the Revert 700 Total Protein Stain, as per the manufacturer’s instructions.

### Real-time quantitative PCR

Total RNA was extracted from HLCs using the Monarch^®^ Total RNA Miniprep Kit (New England BioLabs, T2010). cDNA was generated using the High Capacity cDNA Reverse Transcriptase Kit (Applied Biosystems, 4368814). A master mix was prepared using PerfeCTa FastMix II (Quanta Biosciences, Inc., 95118-250). cDNA was amplified and quantified using the Universal Probe Library (Roche, Burgess Hill, UK) system on a Roche LightCycler 480 (Roche Diagnostics Ltd, Switzerland). Details of primers and Universal Probe Library probes (Roche) and primers used can be found in Table S1.

### Measurement of mitochondrial and nuclear DNA

DNA was extracted from HLCs using the Monarch^®^ Genomic DNA Purification Kit (New England BioLabs, T3010). A master mix was prepared using the Luna® Universal qPCR Master Mix (New England BioLabs, M3003). DNA was amplified and quantified using a Roche LightCycler 480 (Roche Diagnostics Ltd, Switzerland). Primers were designed against two regions of the NCBI mitochondrial reference sequence NC_012920.1, and one region of the genomic reference sequence NC_000011.10. Primers used can be found in Table S2.

### Statistical analyses

All statistical analyses were performed using Graph Prism Version 8.0 for Windows or macOS, GraphPad Software, La Jolla California USA, www.graphpad.com. Normality of data distribution was measured using the Shapiro-Wilks test. Normally distributed data were analysed using a parametric unpaired Student’s t-test, and non-normally distributed data were analysed using a non-parametric Mann-Whitney test. Data were considered to be significant where *p* <0.05.

## Results

### LPO-treatment induces macrovesicular steatosis in HLCs

We previously reported that the high energy substrate cocktail LPO promotes intracellular lipid accumulation in HLCs [17] although it was not known whether LPO promotes lipid droplet biogenesis or increased accumulation of lipids within existing droplets. Stem cell derived HLCs were differentiated and characterised as before [18] (Figure 1A). Analysis of gene expression using RT-qPCR showed increased expression of transcripts for PLIN2, PLIN4, and PLIN5, which are typically associated with intracellular lipid droplet membranes, suggesting an increase in lipid droplet size (Figure 1B-D). Analysis of lipid accumulation using high content imaging (Figure 1E) demonstrated that LPO exposure was not associated with a change in the number of lipid droplets (Figure 1F), but led to a ∼2-fold increase in the size of intracellular lipid droplets (Figure 1G), consistent with the development of macrovesicular steatosis.

**Figure 1.**
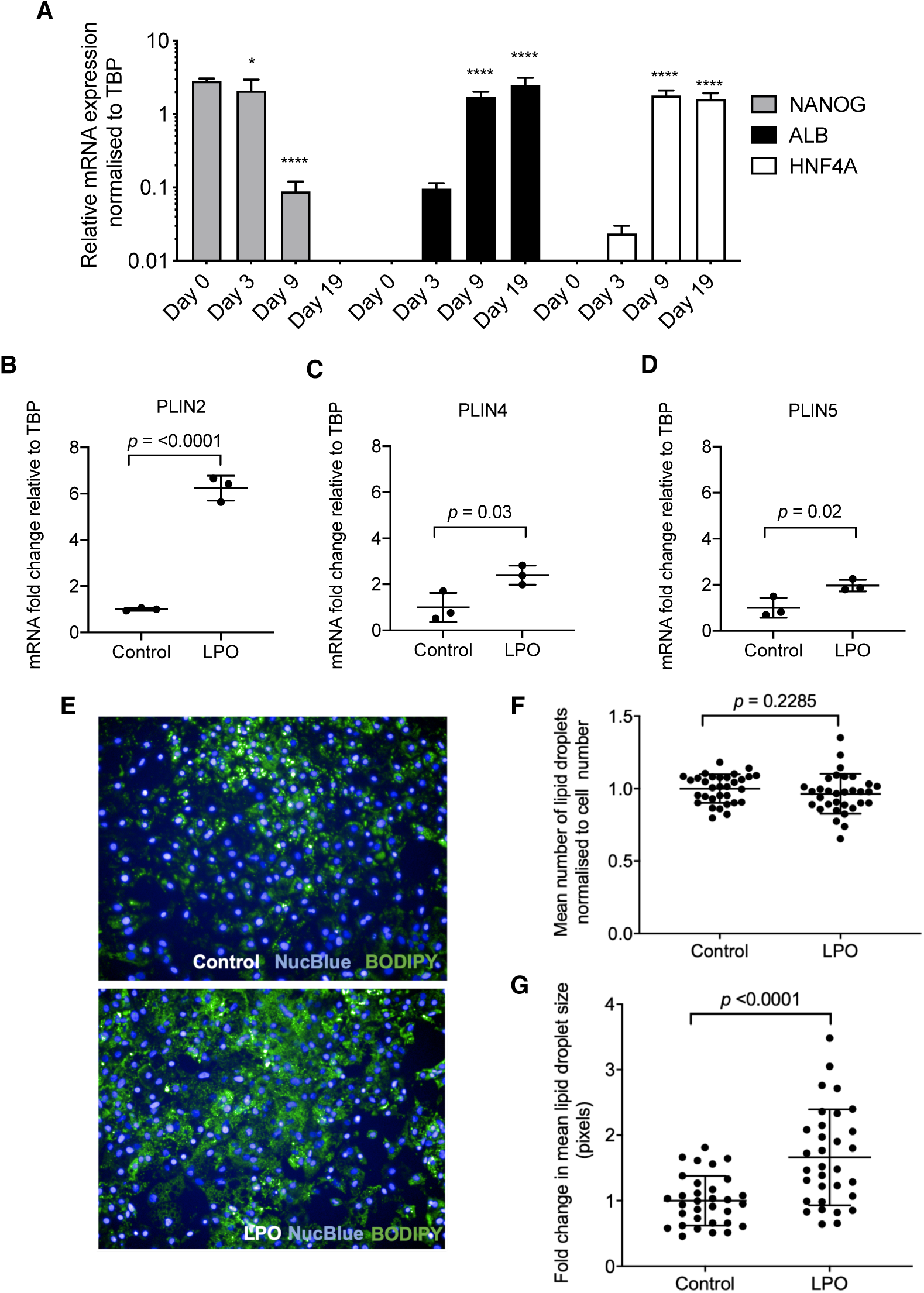
LPO treatment induces development of macrovesicular lipid droplets. (**A**) H9 hPSCs differentiate to HLCs, losing pluripotency and gaining expression of the hepatic markers albumin and HNF4A (n = 3 biological replicates/group); (**B-D**) PLIN2, PLIN4, and PLIN5 expression increases in response to LPO treatment (n = 3 biological replicates per group). (**E**) Representative images of lipid accumulation in control & LPO-treated groups 10x magnification. (**F**) Number of lipid droplets do not increase, (**G**) but intracellular lipid droplets increase in size (n = 32 biological replicates/group). Data analysed using two-tailed Student t-test and expressed as mean ± SD, \**p*<0.05, \*\*\*\**p*<0.0001.

### Macrovesicular steatosis in hepatocytes is associated with electron transport chain dysfunction

It has been suggested that NAFLD-associated macrovesicular steatosis results in impairment of mitochondrial respiration and we therefore proceeded to analyse this in steatotic HLCs (Figure 2A). Basal oxygen consumption rate (OCR), representing combined mitochondrial and non-mitochondrial oxygen consumption, was unchanged following LPO exposure (Figure 2B). Additionally, oligomycin A, a complex V inhibitor decreased OCR equally effectively in control and treatment group. Firstly, this indicated that there were no changes in ATP-linked respiration in response to macrovesicular steatosis (Figure 2C). Secondly, when comparing oligomycin A-induced alterations in OCR with those following addition of rotenone/antimycin A, we could detect no changes in proton leak between groups (Figure 2D). The addition of the ETC uncoupler FCCP revealed a decrease in maximal respiration in the steatotic HLCs, suggesting ETC dysfunction (Figure 2E). Using OCR measurements following FCCP treatment, and comparing to the basal OCR, we calculated that there was a decrease in reserve capacity in the LPO-treated cells, compared with controls (Figure 2F). Subsequently, complex I and III were targeted with rotenone/antimycin A, to completely inhibit oxidative phosphorylation, which reduced OCR to a similar level in both groups, suggesting no difference in non-mitochondrial sources of OCR between the control and steatotic HLCs.

**Figure 2.**
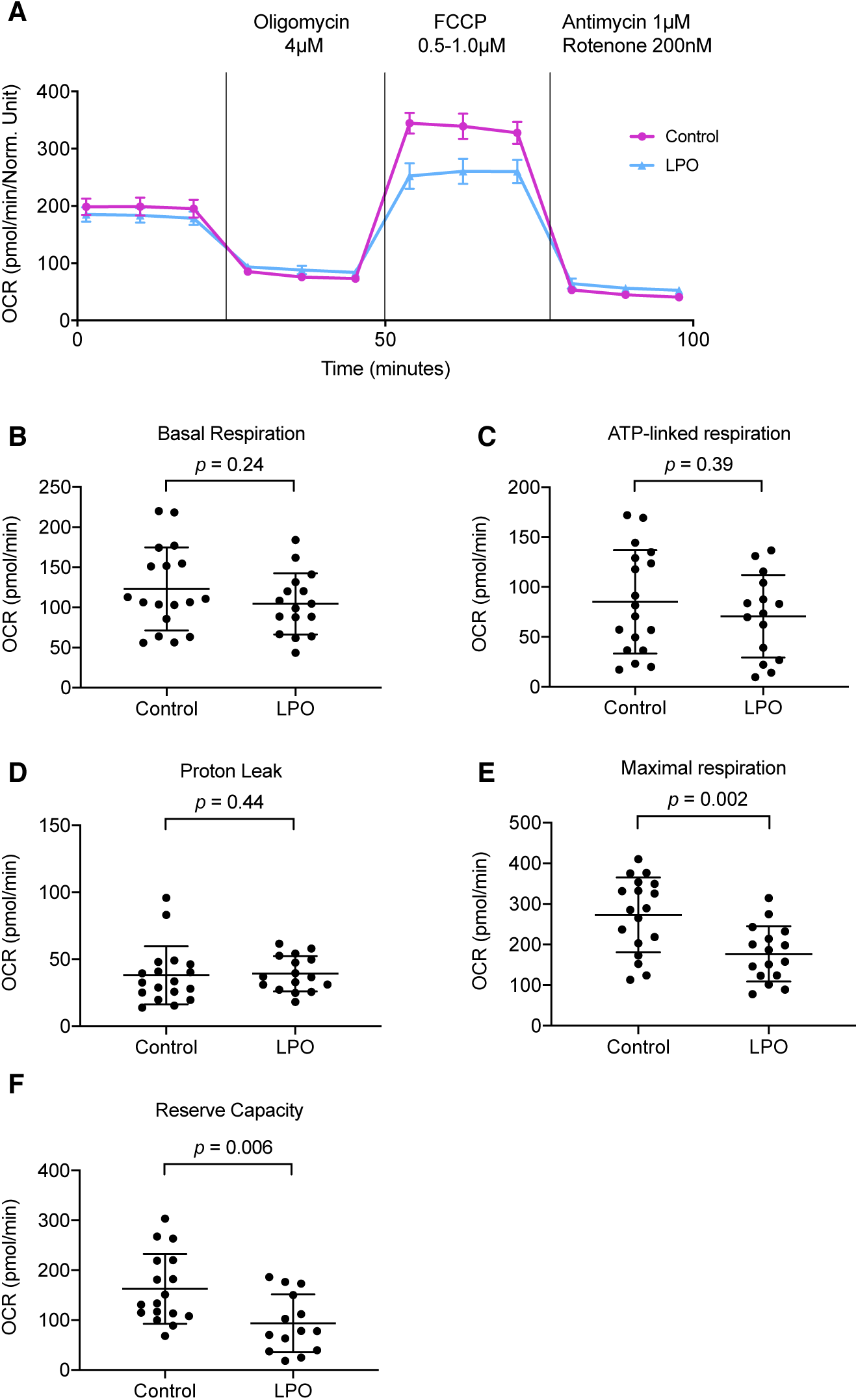
LPO treatment results in decreased maximal respiration in mitochondria but not in other aspects of oxygen consumption rate. HLCs were injected sequentially from ports A-C with 4 μM oligomycin, 0.5-1 μM FCCP, and 1 μM antimycin combined with 200 nM rotenone. (**A**) Raw trace of OCR comparing cells with or without LPO treatment; (**B**) basal respiration; (**C**) ATP-linked respiration; (**D**) proton leak; (**E**) maximal respiration; (**F**) reserve capacity compared to control treatment (*n* = 18 and 16 biological replicates in the control and treatment group, respectively). Data analysed using two-tailed Student t-test for parametric data, or Mann-Whitney U test for non-parametric data, and expressed as mean ± SD.

### Induction of macrovesicular steatosis is not associated with mitochondrial fragmentation or alterations in mitochondrial number

To analyse whether the observed changes in OCR could be due to mitochondrial fragmentation rather than ETC dysfunction we measured citrate synthase (CS) activity. LPO treatment had no impact on CS activity (Figure 3A), suggesting that mitochondria remain intact following intracellular lipid accumulation. A final question was whether changes in ETC function, as demonstrated by decreased cellular maximal respiration and reserve capacity, were due to decreased numbers of mitochondria in response to intracellular lipid accumulation. Since these were the only respiratory measurements to change, we hypothesised that this was not the case. We first measured protein levels of succinate dehydrogenase subunit A (SDHA) and pyruvate dehydrogenase (PDH) (Figure 3B), which localise to the mitochondria. Protein levels of SDHA and PDH did not change in response to intracellular lipid accumulation, suggesting that mitochondrial number is not altered by LPO-induced macrovesicular steatosis. Furthermore, there were no changes in mitochondrial or nuclear DNA (Figure 3C) and no change in the ratio of nuclear to mitochondrial DNA (Figure 3D). Finally, we used a high content microscopy-based approach to measure intracellular mitochondrial quantity (Figure 3E), and observed no changes following exposure to LPO (Figure 3F). Taken together, these data suggest that mitochondrial number is not altered in the presence of intracellular lipid accumulation.

**Figure 3.**
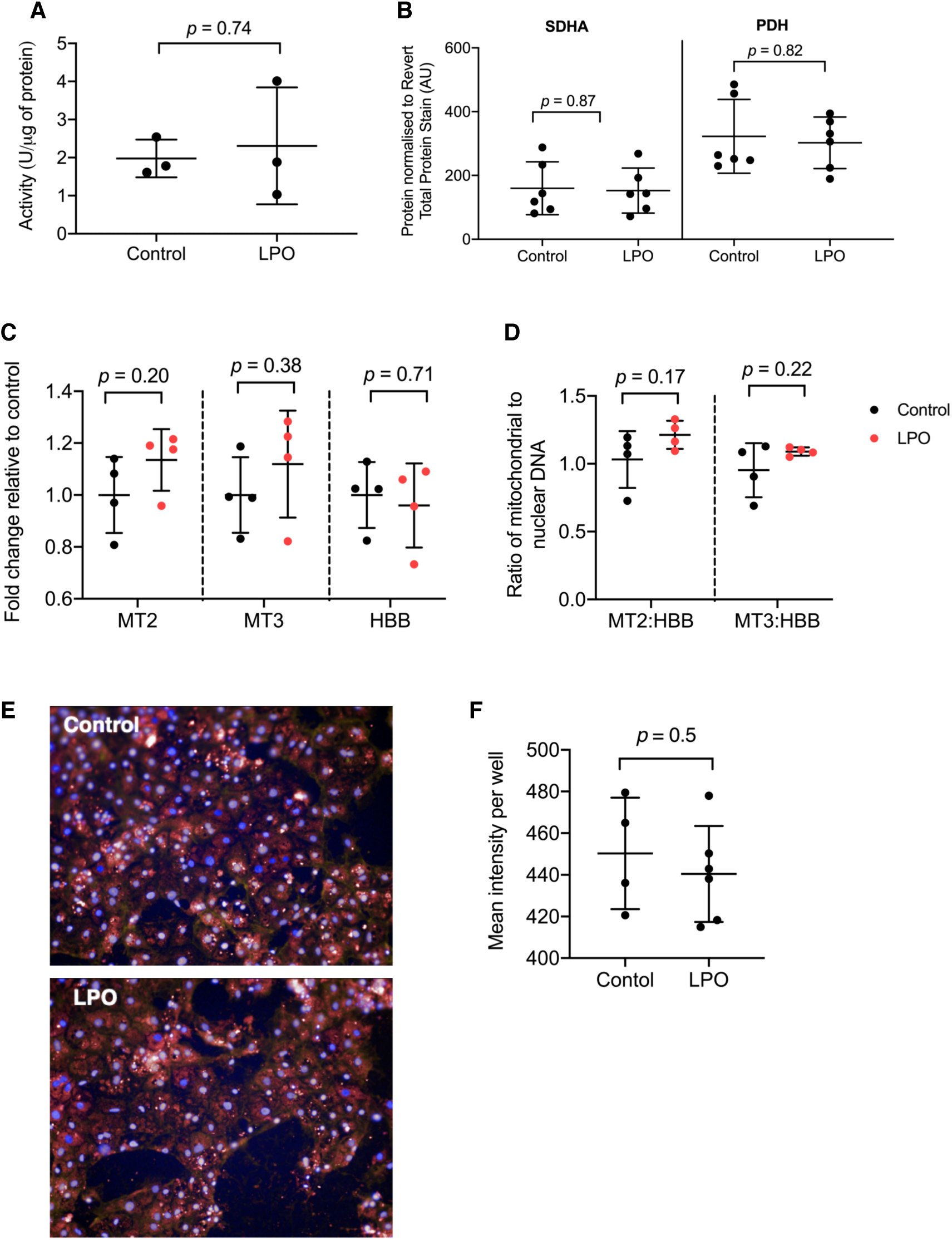
LPO does treatment does induce mitochondrial fragmentation or change mitochondrial number. (**A**) Levels of citrate synthase activity were unaltered by LPO treatment (n = 3 biological replicates/group); (**B**) Protein levels of succinate dehydrogenase subunit A (SDHA) and pyruvate dehydrogenase were unchanged (n = 6 biological replicates/group); (**C**) no changes were observed in abundance of DNA in mitochondrial region 2 (MT2); mitochondrial region 3 (MT3); or genomic region beta-globin (HBB); (**D**) there were no significant changes in the ratio of mitochondrial to genomic DNA, indicating no alterations in the quantity of mitochondria in response to treatment (n=4 biological replicates/group for **C** and **D**); (**E**) Representative images of mitochondrial content in control & LPO-treated groups 10x magnification. Blue staining = NucBlue, red staining = MitoTracker Deep Red; (**F**) high content microscopy revealed no changes in mitochondrial quantity in response to treatment (n = 4 and 6 biological replicates in the control and treatment group, respectively). Data analysed using two-tailed Student t-test for parametric data, or Mann-Whitney U test for non-parametric data, and expressed as mean ± SD.

## Discussion

In this study we aimed to determine the nature of NAFLD-associated changes in mitochondrial respiration, and whether these occur as a result of mitochondrial dysfunction or reductions in mitochondrial number. Using a combination of high content microscopy and gene expression analyses, we first confirmed that treatment of HLCs with LPO induced intracellular lipid accumulation (Figure 1). The observed changes in the expression of genes associated with lipid droplet size (PLIN2, 4 and 5) are also in agreement with findings from human studies with intracellular lipid accumulation associated with an upregulation of PLIN2 in human hepatocytes [21] and biopsies from NAFLD patients [22]. In adipocytes, PLIN4 localises to the surface of unilocular lipid droplets, and may participate in their early development. Although there are no human studies describing perturbed expression of PLIN4 in association with NAFLD, its knockout protects against liver steatosis in mice [23]. Furthermore, PLIN4 knockout diminishes the accumulation of triacylglycerol in lipid droplets and reduces the expression of PLIN5 in mouse cardiac tissue. Therefore, their interplay may be important for the pathology of NAFLD. In support of this, *in vitro* studies show that PLIN5 blocks lipid droplet lipolysis in hepatocytes and an increase in PLIN5 expression may therefore contribute to the development of macrovesicular steatosis [24]. Increases in the expression of PLIN2, PLIN4 and PLIN5 in LPO-treated HLCs, alongside BODIPY staining, strongly indicate that this model recapitulates macrovesicular steatosis observed in humans with NAFLD.

Following the further establishment of relevance to human disease, we wished to understand whether macrovesicular steatosis was linked to dysfunction of mitochondrial respiration [15]. Previous studies have shown that mitochondrial maximal respiration increases in obese humans with NAFLD but decreases on progression to NASH, despite an increase in mitochondrial mass. In contrast to the initial hypothesis, we observed decreased maximal respiration and reserve capacity in cells displaying a NAFLD phenotype. This indicates that whilst mitochondria are mostly able to maintain respiratory integrity in LPO-exposed cells, the ETC, and therefore oxidative phosphorylation, may be compromised. Given that these were the only parameters of mitochondrial respiration to change, it suggests that LPO treatment of HLCs inhibits complexes I-III of the ETC. This effect may arise due to the use of octanoic acid, which inhibits complex I-III in rat liver, as well as increasing oxidative stress through reactive oxygen species production [25]. Inhibition of complex II may also impact on the TCA cycle, potentially altering its function, and the generation of cofactors necessary for oxidative phosphorylation, further impacting on cellular metabolism. However, further studies are required to test these hypotheses.

Finally, we questioned whether changes in maximal respiration arose from fragmentation of mitochondria or resulted from increased biogenesis. Given that maximal respiration was the only parameter of respiration that changed, we suspected this was not the case. This was confirmed by lack of change in the ratio of nuclear to mitochondrial DNA, suggesting that mitochondrial biogenesis was not affected by LPO treatment in these studies. This is consistent with human studies of NAFLD which show no changes in mitochondrial mass in steatotic versus non-steatotic liver tissue [15]. In contrast, liver tissue from patients with NASH contains greater numbers of mitochondria, with a concomitant decrease in maximal respiration. Therefore, the model of NAFLD presented here may mirror the transition between NAFLD and NASH, with respiratory dysfunction in the absence of mitochondrial biogenesis. This discrepancy may also be due to previous studies using isolated mitochondria from whole tissues, comprised of multiple cell types, whereas we examined a single cell population.

In conclusion, here we demonstrate that the treatment of HLCs with LPO induces intracellular lipid accumulation in patterns similar to those observed in human pathology. Furthermore, we observed decreased mitochondrial maximal respiration, in the absence of altered mitochondrial biogenesis. Taken together, our data demonstrate that hepatocyte intracellular lipid accumulation is linked to mitochondrial dysfunction in response to the changing bioenergetic demands associated with NAFLD, impacting on the ETC, and limiting maximal respiration.

## Acknowledgements

MCS was supported by a British Heart Foundation PhD studentship (FS/16/54/32730) and by the BHF Centre of Research Excellence. BLV and DCH were supported by an award from the Chief Scientist Office (TCS/16/37). JMR was supported by an MRC PhD studentship. MJL was supported by a Wellcome Trust PhD studentship as part of the Edinburgh Clinical Academic Track (102839/Z/13/Z). RNC and NMM were supported by a Wellcome Trust New Investigator Award to NMM (100981/Z/13/Z). AJD was supported by the BHF Centre of Research Excellence, Edinburgh. We thank Will Cawthorn for discussions about mitochondrial quantification.

## Author contributions

MCS, JMR, BLV, MJL, RNC and NMM performed experiments. DCH and AJD conceived the experiments. MCS, NMM, DCH and AJD wrote the paper. All authors contributed to drafts of the paper and approved the final version.

## Supplementary materials S1

Primer sequences and UPL probes used for qPCR assays

## References

1. World Health Organization. Obesity and overweight. https://www.who.int/news-room/fact-sheets/detail/obesity-and-overweight (2018).

2. Valenti, L., Bugianesi, E., Pajvani, U. & Targher, G. Nonalcoholic fatty liver disease: cause or consequence of type 2 diabetes? Liver International vol. 36 1563–1579 (2016).

3. Wang, L. & Yu, S. Pathology of non-alcoholic fatty liver disease. Int. J. Dig. Dis. 2, (2016).

4. Asrih, M. & Jornayvaz, F. R. Metabolic syndrome and nonalcoholic fatty liver disease: Is insulin resistance the link? Molecular and Cellular Endocrinology vol. 418 55–65 (2015).

5. Laursen, T. L. et al. Bariatric surgery in patients with non-alcoholic fatty liver disease - From pathophysiology to clinical effects. World Journal of Hepatology vol. 11 138–249 (2019).

6. Sunny, N. E., Parks, E. J., Browning, J. D. & Burgess, S. C. Excessive hepatic mitochondrial TCA cycle and gluconeogenesis in humans with nonalcoholic fatty liver disease. Cell Metab. 14, 804–810 (2011).

7. Satapati, S. et al. Elevated TCA cycle function in the pathology of diet-induced hepatic insulin resistance and fatty liver. J. Lipid Res. 53, 1080–1092 (2012).

8. Patterson, R. E. et al. Lipotoxicity in steatohepatitis occurs despite an increase in tricarboxylic acid cycle activity. Am. J. Physiol. - Endocrinol. Metab. 310, ajpendo.00492.2015 (2016).

9. Sunny, N. E., Bril, F. & Cusi, K. Mitochondrial adaptation in nonalcoholic fatty liver disease: novel mechanisms and treatment strategies. Trends Endocrinol. Metab. 28, 250–260 (2017).

10. Rector, R. S. et al. Mitochondrial dysfunction precedes insulin resistance and hepatic steatosis and contributes to the natural history of non-alcoholic fatty liver disease in an obese rodent model. J. Hepatol. 52, 727–736 (2010).

11. Sanyal, A. J. et al. Nonalcoholic steatohepatitis: association of insulin resistance and mitochondrial abnormalities. Gastroenterology 120, 1183–1192 (2001).

12. Caldwell, S. H. et al. Mitochondrial abnormalities in non-alcoholic steatohepatitis. J. Hepatol. 31, 430–434 (1999).

13. Fabbrini, E. et al. Alterations in adipose tissue and hepatic lipid kinetics in obese men and women with nonalcoholic fatty liver disease. Gastroenterology 134, 424–431 (2008).

14. Satapati, S. et al. Mitochondrial metabolism mediates oxidative stress and inflammation in fatty liver. J. Clin. Invest. 125, 4447–4462 (2015).

15. Koliaki, C. et al. Adaptation of hepatic mitochondrial function in humans with non-alcoholic fatty liver is lost in steatohepatitis. Cell Metab. 21, 739–746 (2015).

16. Knott, A. B., Perkins, G., Schwarzenbacher, R. & Bossy-Wetzel, E. Mitochondrial fragmentation in neurodegeneration. Nature Reviews Neuroscience vol. 9 505–518 (2008).

17. Lyall, M. J. et al. Modelling non-alcoholic fatty liver disease in human hepatocyte-like cells. Philos. Trans. R. Soc. B Biol. Sci. 373, 20170362 (2018).

18. Wang, Y. et al. Defined and scalable generation of hepatocyte-like cells from human pluripotent stem cells. J. Vis. Exp. e55355–e55355 (2017).

19. Bray, M. A. et al. Cell Painting, a high-content image-based assay for morphological profiling using multiplexed fluorescent dyes. Nat. Protoc. 11, 1757–1774 (2016).

20. Orellana, E. & Kasinski, A. Sulforhodamine B (SRB) assay in cell culture to investigate cell proliferation. Bio-Protocol 6, (2016).

21. Sahini, N. & Borlak, J. Genomics of human fatty liver disease reveal mechanistically linked lipid droplet–associated gene regulations in bland steatosis and nonalcoholic steatohepatitis. Transl. Res. 177, 41–69 (2016).

22. Fujii, H. et al. Expression of perilipin and adipophilin in nonalcoholic fatty liver disease; relevance to oxidative injury and hepatocyte ballooning. J. Atheroscler. Thromb. 16, 893–901 (2009).

23. Chen, W. et al. Inactivation of Plin4 downregulates Plin5 and reduces cardiac lipid accumulation in mice. Am. J. Physiol. - Endocrinol. Metab. 304, E770–9 (2013).

24. Wang, C. et al. Perilipin 5 improves hepatic lipotoxicity by inhibiting lipolysis. Hepatology 61, 870–882 (2015).

25. Scaini, G. et al. Toxicity of octanoate and decanoate in rat peripheral tissues: evidence of bioenergetic dysfunction and oxidative damage induction in liver and skeletal muscle. Mol. Cell. Biochem. 361, 329–335 (2012).

