## Supplemental Tables for "Nonalcoholic fatty liver disease is associated with decreased hepatocyte mitochondrial respiration, but not mitochondrial number" (from another version of this preprint: version 1, 2020-03-11)

| **Gene** | **Forward Primer** | **Reverse Primer** | **Probe #** |
| --- | --- | --- | --- |
| NANOG | ATGCCTCACACGGAGACTGT | CAGGGCTGTCCTGAATAAGC | 69 |
| ALB | GTGAGGTTGCTCATCGGTTT | GAGCAAAGGCAATCAACACC | 7 |
| HNF4A | AGCAACGGACAGATGTGTGA | TCAGACCCTGAGCCACCT | 27 |
| PLIN2 | TCAGCTCCATTCTACTGTTCACC | CCTGAATTTTCTGATTGGCACT | 72 |
| PLIN4 | AGTTCCAAGCCAGGGACAC | TGCTGGGCCTTTTCAATC | 1 |
| PLIN5 | TACAGTGCAGCCAAGGACAG | CGCACACGCAGTTCTCAG | 3 |

**Table S1.** Primer pairs and probes used to quantify mRNA expression

| **Gene** | **Forward Primer** | **Reverse Primer** |
| --- | --- | --- |
| MT1 | CTCACTCTCACTGCCCAAGA | TGAGAATGAGTGTGAGGCGT |
| MT2 | ACCCACCAATCACATGCCTA | GTGTTACATCGCGCCATCAT |
| HBB | TGGTGCATCTGACTCCTGAG | TCTCCACATGCCCAGTTTCT |

**Table S2.** Primer pairs used to quantify mitochondrial and nuclear DNA
